## Supplementary figures and images for "Natural variation in expression of a plant immune receptor mediates elicitor sensitivity"

### S1 Figure

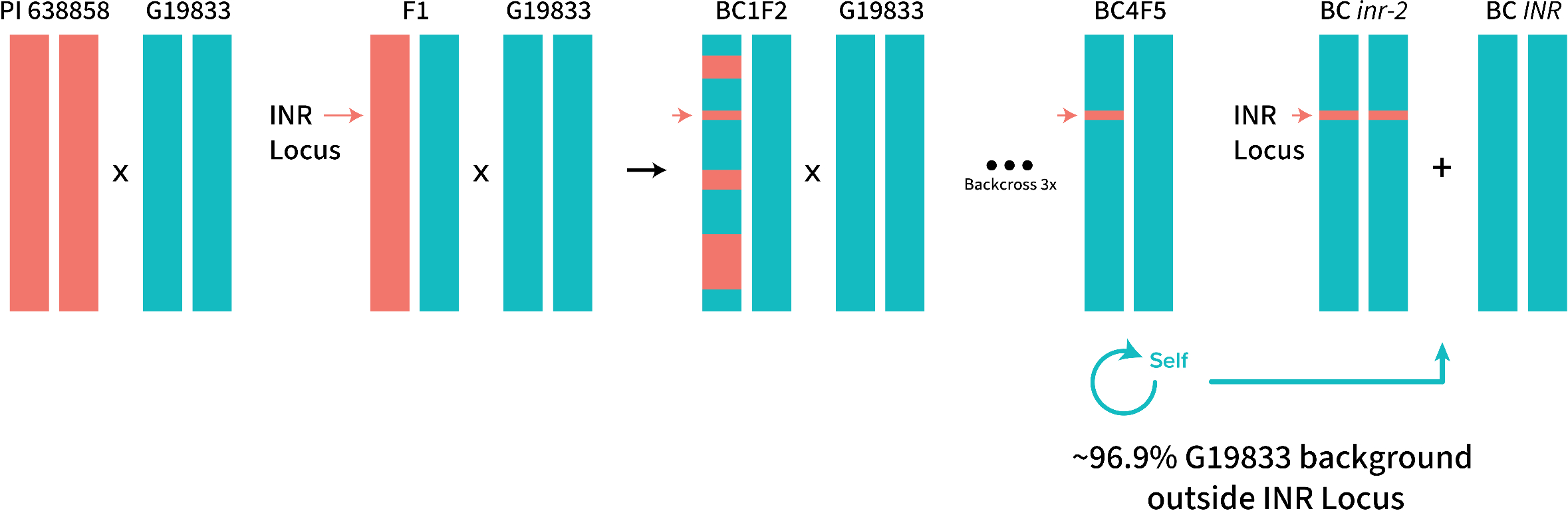

### S2 Figure

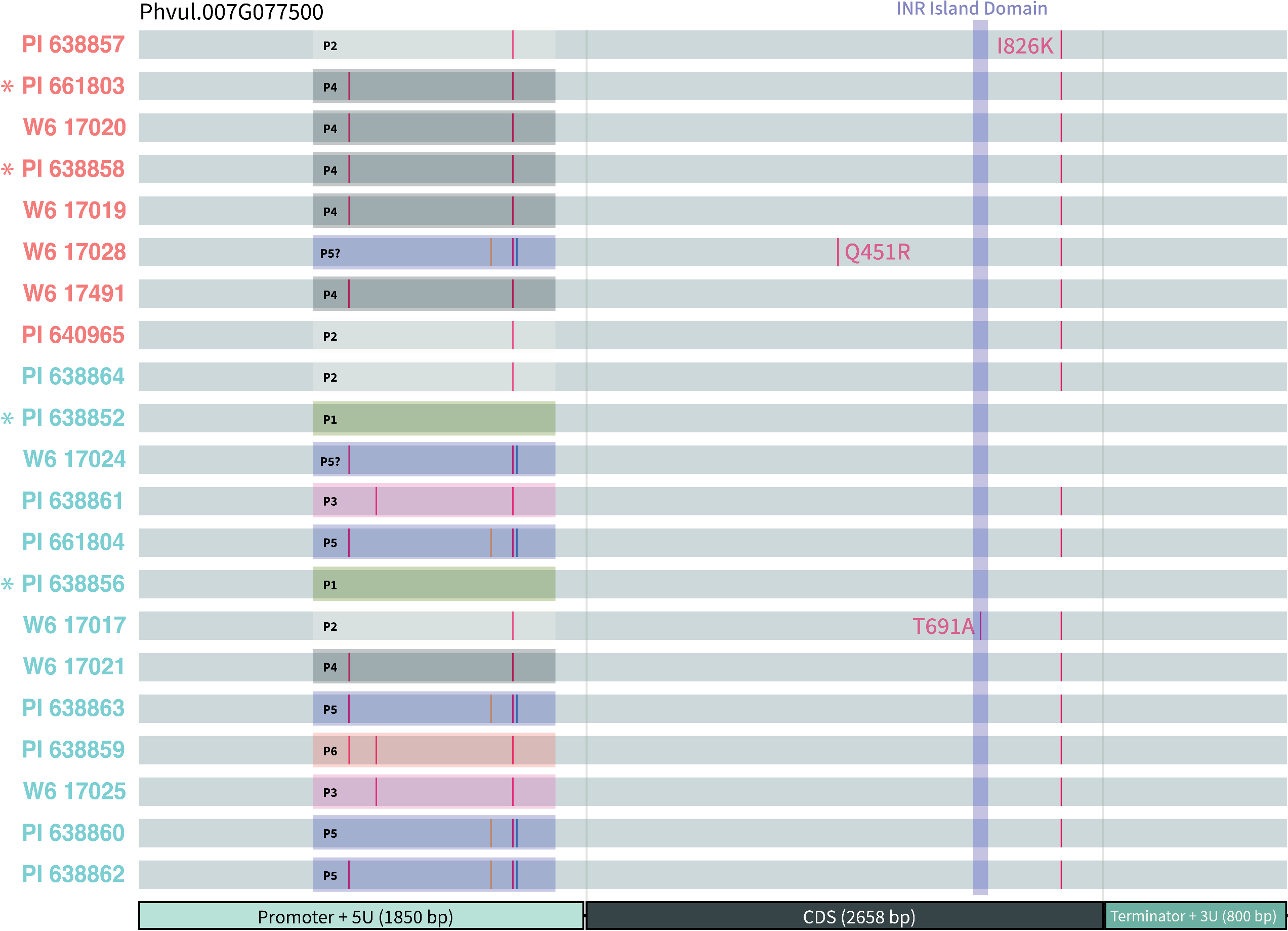

### S3 Figure

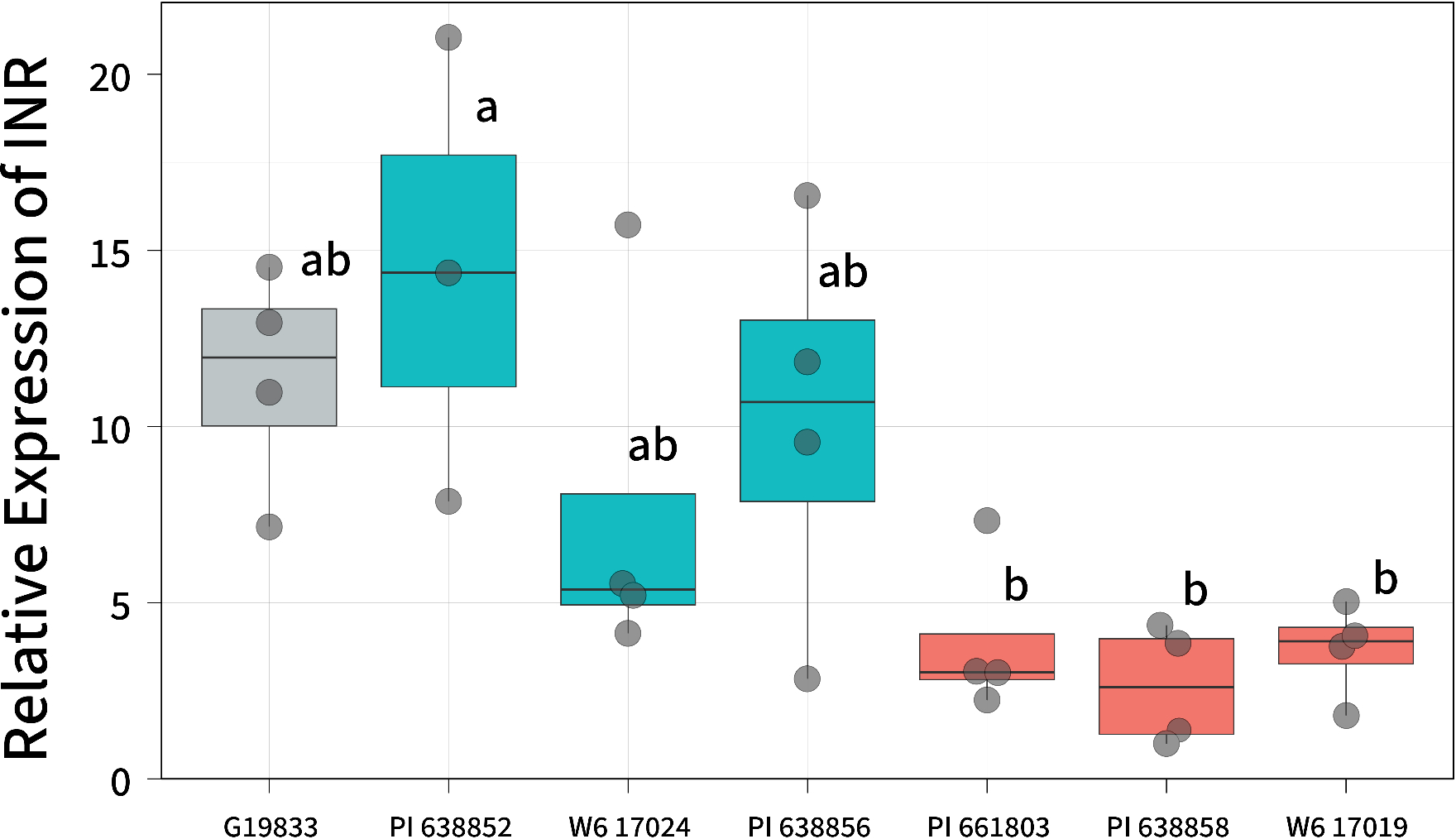

### S4 Figure

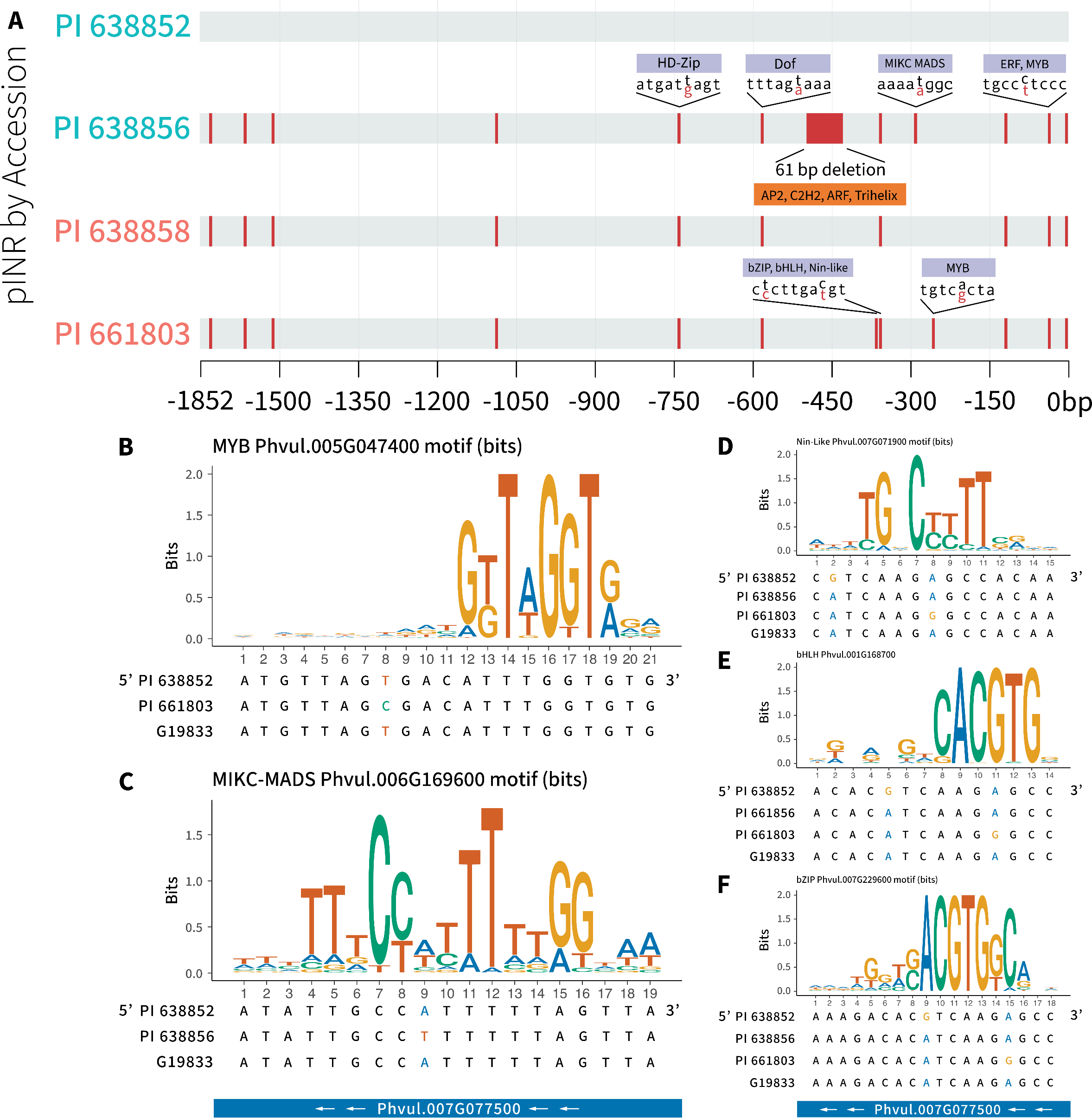

### S5 Figure

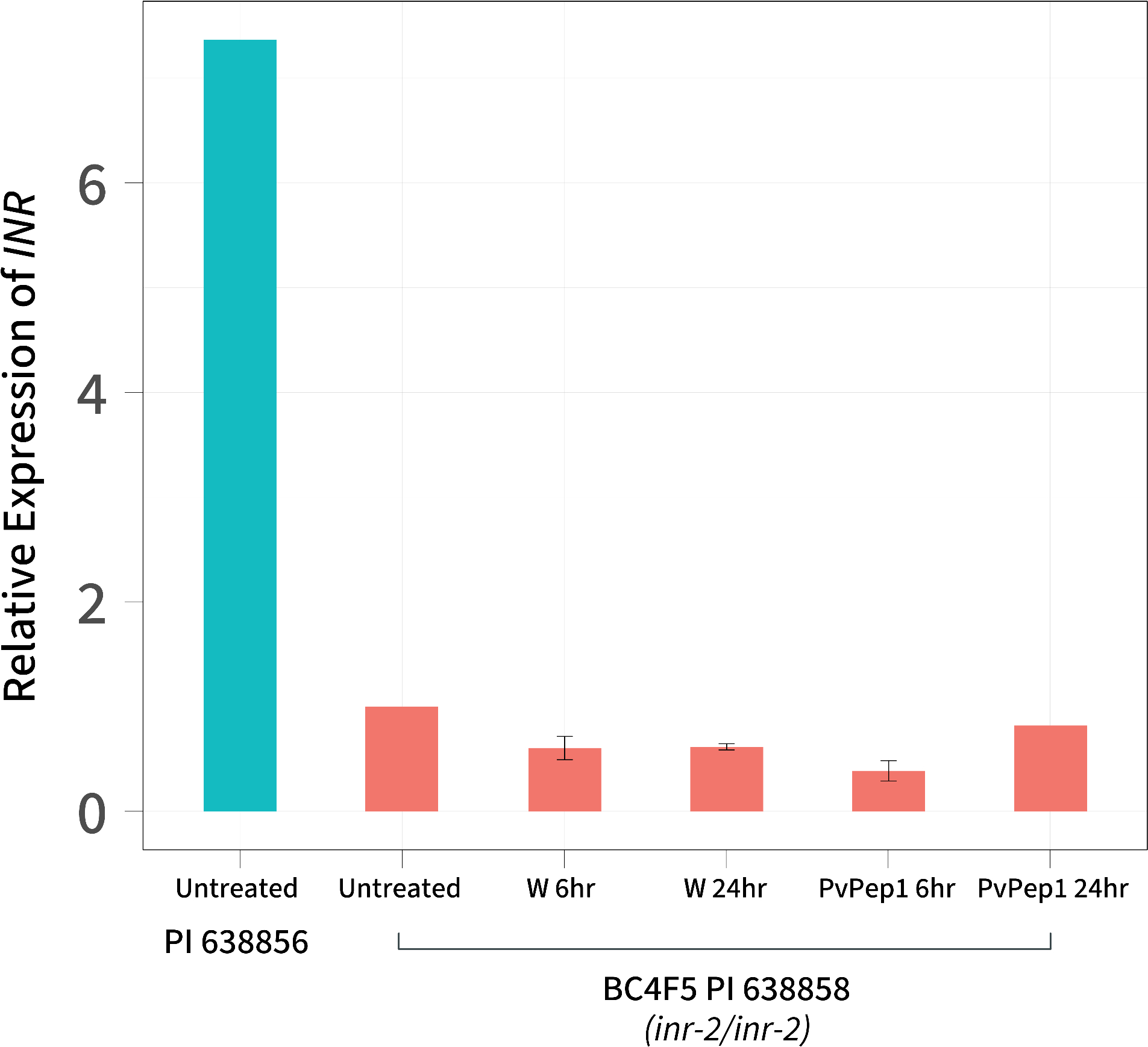

### S6 Figure

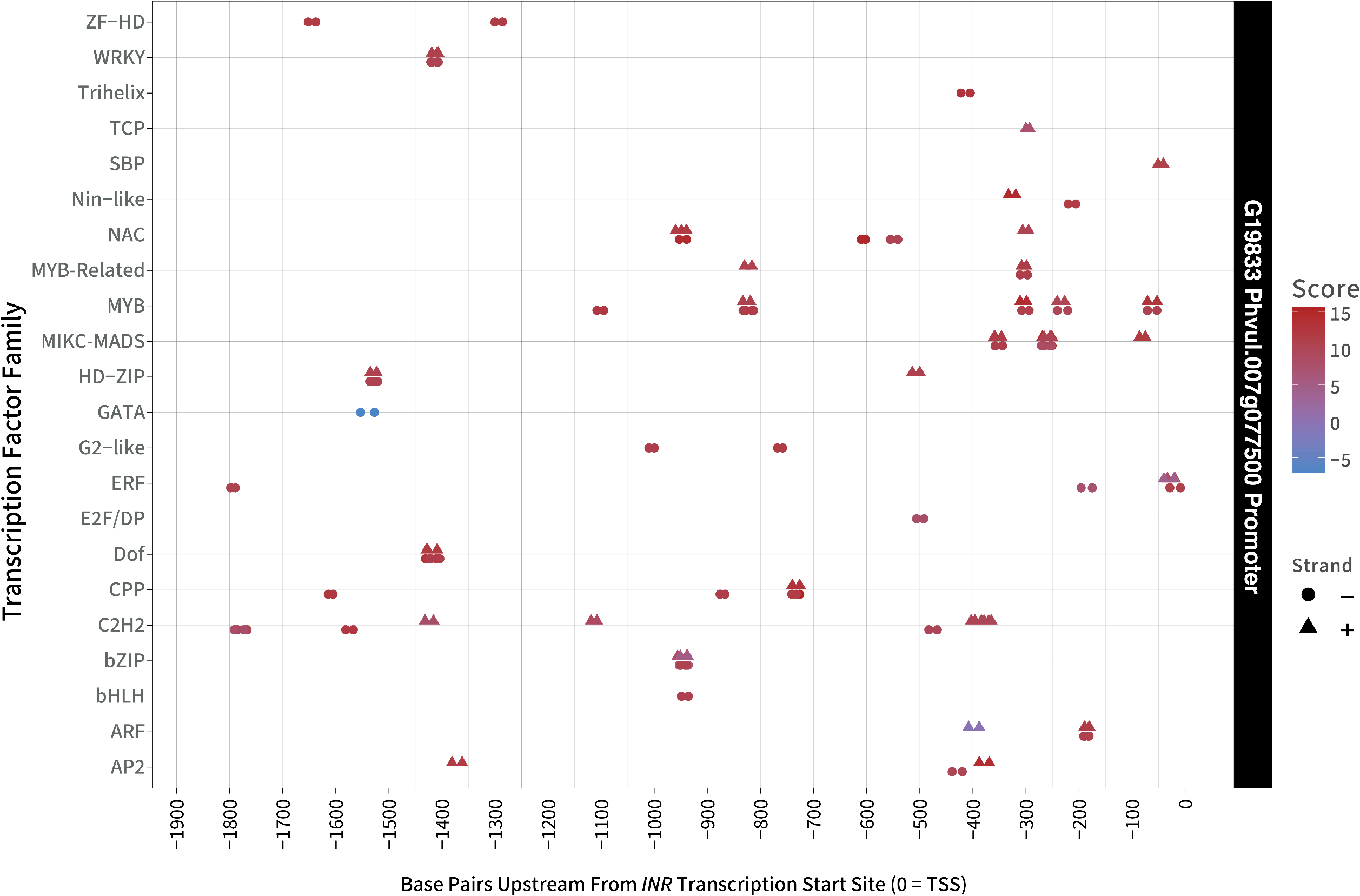

### S7 Figure

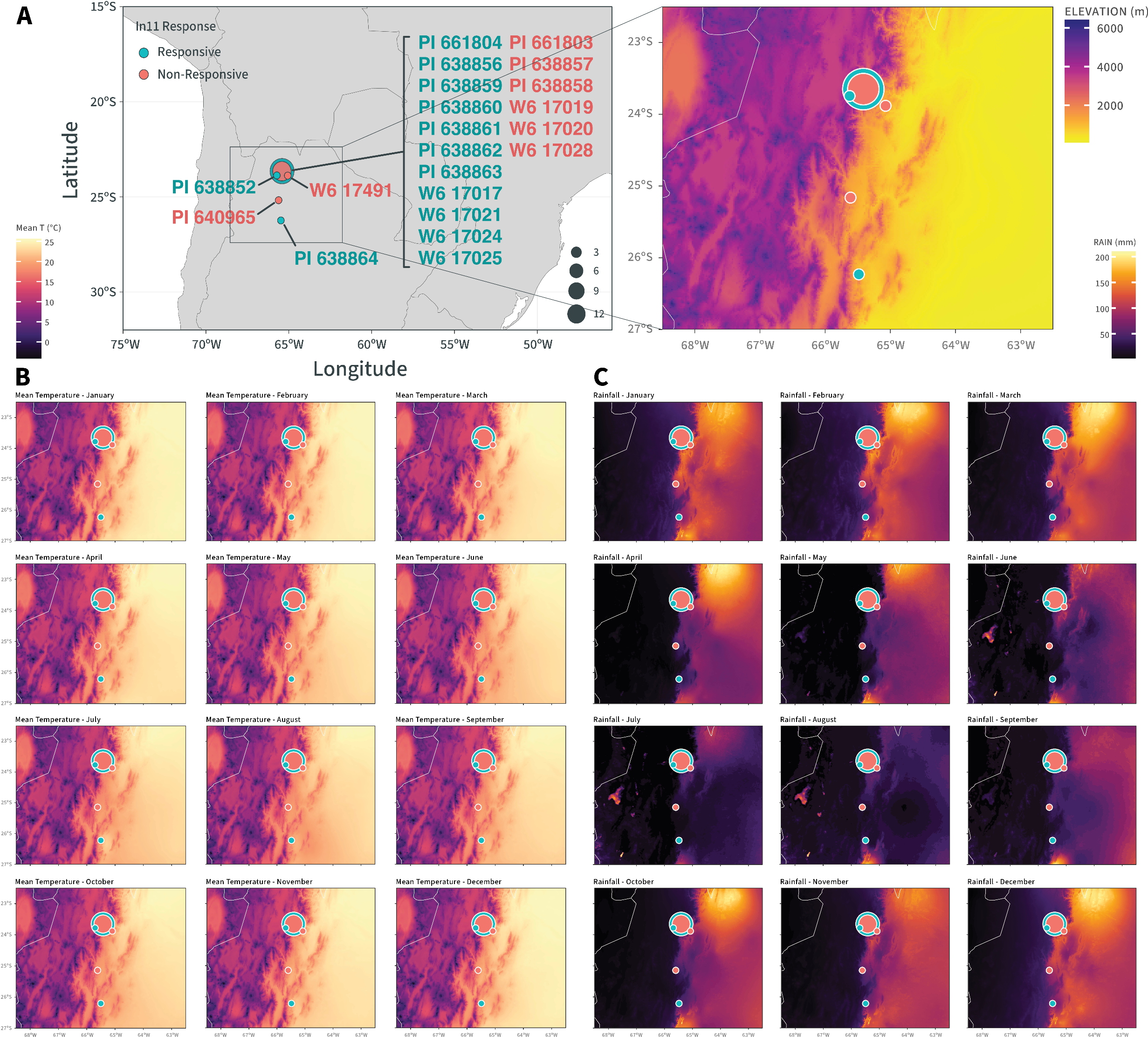
